# Circular RNA expression and regulatory network prediction in posterior cingulate astrocytes in elderly subjects

**DOI:** 10.1101/268888

**Authors:** Shobana Sekar, Lori Cuyugan, Jonathan Adkins, Philipp Geiger, Winnie S. Liang

**Affiliations:** Neurogenomics division, Translational Genomics Research Institute, Phoenix, AZ, 85004, USA; Arizona Alzheimer’s Consortium, Phoenix, AZ, 85014, USA; Department of Biomedical Informatics, Arizona State University, Tempe, AZ, 85287, USA

**Keywords:** Circular RNAs, astrocytes, posterior cingulate, aging, regulatory network

## Abstract

**Background:** Circular RNAs (circRNAs) are a novel class of endogenous, non-coding RNAs that form covalently closed continuous loops and are both highly conserved and abundant in the mammalian brain. A role for circRNAs in sponging microRNAs (miRNAs) has been proposed, but the circRNA-miRNA-mRNA interaction networks in human brain cells have not been defined. Therefore, we identified circRNAs in RNA sequencing data previously generated from astrocytes microdissected from the posterior cingulate (PC) of Alzheimer’s disease (AD) patients (N=10) and healthy elderly controls (N=10) using four circRNA prediction algorithms - CIRI, CIRCexplorer, find_circ and KNIFE.

**Results:** Overall, utilizing these four tools, we identified a union of 4,438 unique circRNAs across all samples, of which 70.3% were derived from exonic regions. Notably, the widely reported CDR1as circRNA was detected in all samples across both groups by find_circ. Given the putative miRNA regulatory function of circRNAs, we identified potential miRNA targets of circRNAs, and further, delineated circRNA-miRNA-mRNA networks using in silico methods. Pathway analysis of the genes regulated by these miRNAs identified significantly enriched immune response pathways, which is consistent with the known function of astrocytes as immune sensors in the brain.

**Conclusions:** In this study, we performed circRNA detection on cell-specific transcriptomic data and identified potential circRNA-miRNA-mRNA regulatory networks in PC astrocytes. Given the known function of astrocytes in cerebral innate immunity and our identification of significantly enriched immune response pathways, the circRNAs we identified may be associated with such key functions. While we did not detect recurrent differentially expressed circRNAs in the context of healthy controls or Alzheimer’s, we report for the first time circRNAs and their potential regulatory impact in a cell-specific and region-specific manner in aged subjects. These predicted regulatory network and pathway analyses may help provide new insights into transcriptional regulation in the brain.

## 1. Background

CircRNAs are a class of endogenous, non-coding RNAs that form covalently closed continuous loops and are pervasively expressed in eukaryotes [1–3]. Though RNA circularization events were reported in the 1970s and 1990s [4–6], they were disregarded as molecular artifacts arising from aberrant splicing. However, with the advent of next-generation sequencing technology, coupled with the development of computational algorithms to specifically detect these back-splicing events, numerous circRNAs have been reported since 2012. CircRNAs exhibit cell type-, tissue- and developmental stage-specific expression [3, 7], and show evolutionary conservation between mouse and human [2, 3]. Furthermore, circRNAs are highly abundant in the mammalian brain compared to other tissues such as lungs, heart, kidney, testis and spleen in humans as well as in mouse neuronal cell lines [8], and are derived preferentially from neural genes [9].

The abundance and evolutionary conservation of circRNAs suggests that they could play important roles in cellular processes. A few possible functions have been reported, including microRNA (miRNA) sponges [3, 6, 10, 11], mediation of protein-protein interactions [12] and regulation of parental gene transcription [13]. Furthermore, a few circRNAs have been found to originate from disease-associated genomic loci, suggesting that circRNAs may regulate pathological processes [14–18]. Given these data, it is likely that circRNAs regulate RNA and protein networks, especially in the brain, but the regulatory pathways are still unknown.

In the present study, we characterized the expression and abundance of circRNAs in next generation RNA-sequencing (RNAseq) data of human brain astrocytes. Astrocytes, the most abundant glial cells, play several essential roles in the central nervous system, including homeostasis [19], immunity [20] and energy storage and metabolism [21, 22]. We previously evaluated these astrocytes, which were derived from the posterior cingulate (PC) of Alzheimer’s disease (AD) and healthy elderly control brains (age > 65), and identified AD-associated gene expression changes [23]. For this study, we used four circRNA prediction algorithms to identify circRNAs in these AD and control samples. Given the potential miRNA regulatory function of circRNAs, we then performed in silico identification of miRNA binding sites on the detected circRNAs, and further delineated putative circRNA-miRNA-mRNA networks in astrocytes. We describe here the first astrocyte-specific characterization of circRNAs and their interaction networks in elderly individuals.

## 2. Results

### 2.1 CircRNA detection in PC astrocytes

The RNAseq data generated from our previous study was used for analysis [23]. This data set was generated from 20 human PC astrocyte pools: 10 from late-onset AD (LOAD) brains and 10 from no disease (ND) healthy elderly control brains. Over 85,000,000 reads were sequenced for each sample, with an average mapping percentage of 70.8. On the FASTQ files generated from sequencing, we ran four circRNA prediction algorithms - CIRCexplorer [24], CIRI [25], find_circ [3], and KNIFE [26], and detected a total of 4,438 unique circRNAs with at least two supporting junction reads (Additional file 1: Table S1). Among the detected circRNA candidates, a total of 2,331 circRNAs were identified in the AD samples and 2,425 in the ND samples by at least one of the algorithms (Figure 1a). While 80% of the detected circRNAs had less than ten supporting reads (Figure 1b), 43 circRNAs had over 20 junction reads and were detected in more than one sample, and 31 circRNA candidates were detected in at least five samples with five or more supporting reads. Notably, the widely reported CDR1as circRNA was detected with a median read count of 52, by find_circ in all 20 samples and by CIRI in one of the samples. CircRNA 2:40655612-40657444 (chromosome:start-end) was detected in 12 of the 20 samples by two, three or all four algorithms in each sample (Additional file 1: Table S1). Furthermore, 548 circRNAs detected in our dataset were also reported in the four studies deposited in circBase [27] (Additional file 1: Table S1); various cell lines and tissue types were evaluated in these studies, including cerebellum, diencephalon, SH-SY5Y cells, Hs68 cells, HeLa cells and HEK293 cells.

**Figure 1:**
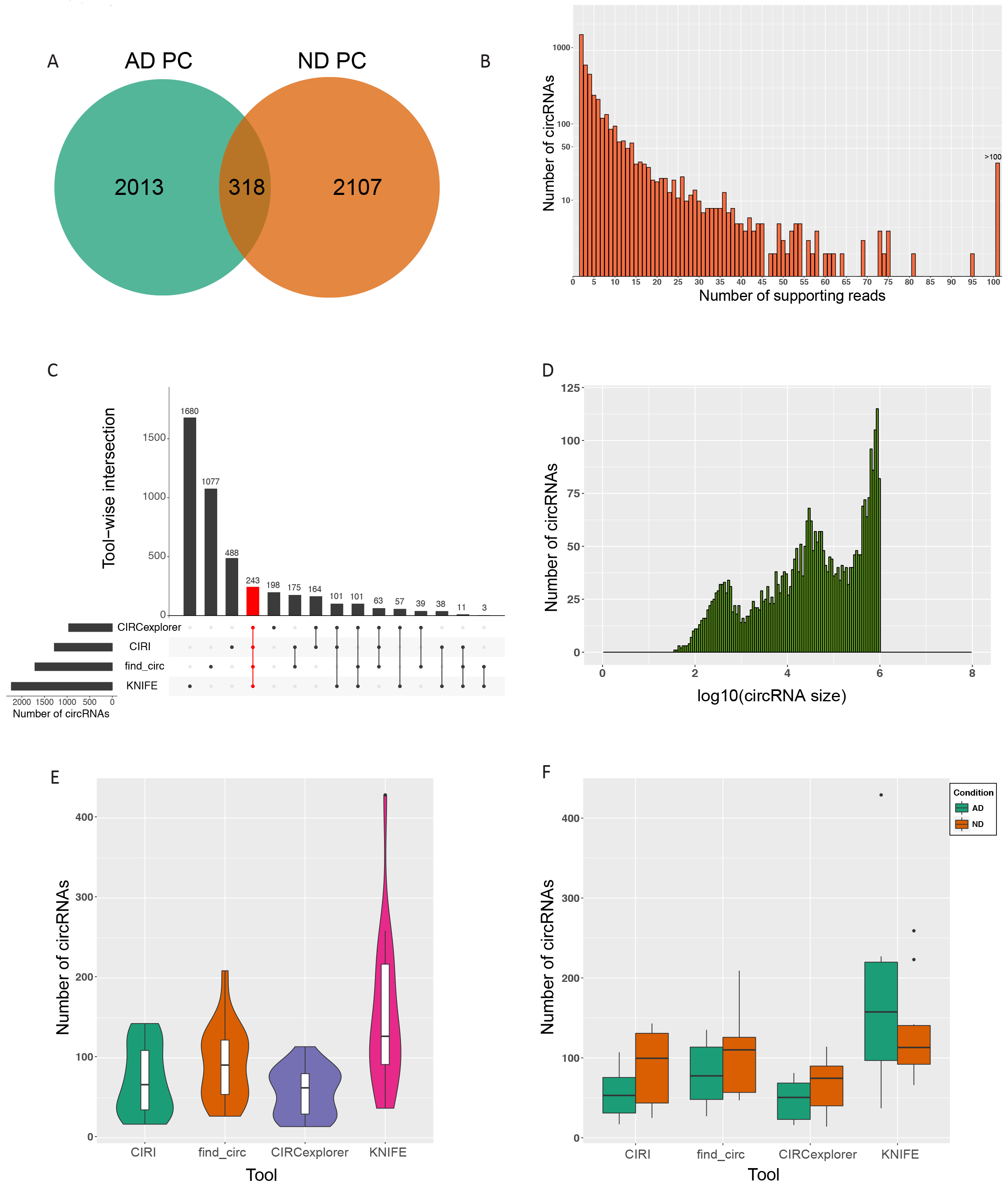
Summary of circRNA prediction results. (a) Number of unique and common circRNAs in AD and ND PC. (b) Read count distribution of all detected circRNAs. (c) Intersection of circRNAs called by the four tools; the red bar indicates the number of circRNAs called by all four tools (d) Size distribution of all detected circRNAs. (e) Violin plots indicating the number of circRNAs predicted by each tool across PC samples along with the probability density. (f) Number of circRNAs predicted by each tool across PC samples, condition-wise. AD, Alzheimer’s disease; ND, no disease; circRNA, circular RNA; PC, posterior cingulate; bp, base pairs.

Among all identified circRNAs, 416 were on chromosome 1 (length = 249,250,621 base pairs), while only eight were detected on chromosome Y (length = 59,373,566 base pairs), consistent with previous findings that the number of circRNAs detected is proportional to the length of the chromosome [28]. Based on RefSeq annotations, we observed that 70.3% of our candidates were derived from exonic regions (3,123/4,438), of which 94% (2,936/3,123) were in coding DNA sequences (CDS; excludes untranslated regions) (Additional file 1: Table S1). Among the exonic circRNAs, 56.4% spanned one to 15 exons per circRNA, of which 20% were derived from single exons, while a small percentage of the exonic circRNAs (6.8%) spanned over 100 exons per circRNA.

As previously reported [29], we observed that the overlap among the circRNAs detected by the different tools was low. Overall, 243 circRNAs were predicted by all four tools, while each tool also predicted unique circRNAs (KNIFE—1680, find_circ—1077, CIRI—488, CIRCexplorer—198; Figure 1c). Most of the candidates called by all the tools originated from CDS (242/243; 99.5%) as well as intronic regions (232/243; 95.5%), and 75% of the exonic candidates spanned two to six exons per circRNA. Further, the size distribution of all detected circRNAs, and the tool-wise and condition-wise distribution of the circRNAs, are summarized in Figures 1d, e and f.

We next compared the relative abundance of circRNAs and corresponding linear RNAs using back-spliced reads and linearly spliced reads with the same splice sites (Methods; Table S2; Figure S1). We observed that for 26 circRNAs, the circular-to-linear ratio was 10 or greater and the linear count was not0, such as circRNA 17:48823196-48824063 from *LUC7L3* (LUC7 like 3 pre-mRNA splicing factor; average back-spliced reads: 413, average linear reads: 16.32) and 1:67356836-67371058 from *WDR78* (WD repeat domain 78; average back-spliced reads: 116.50, average linear reads: 9.50). Further, 44.6% (1,983/4,438) had no expression of linear RNA and 45.5% (2,018/4,438) had higher expression of the linear RNA.

### 2.2 miRNA target prediction and delineation of circRNA-miRNA-mRNA regulatory networks

Given the potential miRNA regulatory function of circRNAs, we next used the miRNA target prediction algorithms miRanda [30] and RNAHybrid [31] to predict the miRNA targets of the circRNAs detected in ten or more samples by at least one of the circRNA prediction algorithms (N = 10 circRNA candidates). Using a list of 2,588 published miRNAs from miRBase [32], we detected 14,296 unique interactions between circRNAs and miRNAs that were predicted by both the miRNA target prediction algorithms and having a miRanda match score >=150. These interactions represent binding sites for miRNAs on each circRNA candidate, predicted based on complementarity in the miRNA seed region (nucleotide positions 2-7 in the miRNA 5’-end). 2,398 miRNAs in the reference set were predicted to have binding sites on our input list of circRNAs. Among these, a set of 612 circRNA-miRNA interaction pairs were predicted to contain over 100 putative interaction sites by the miRanda algorithm (Additional file 3: Table S3). These 612 circRNA-miRNA interactions were predicted for six unique circRNAs and 448 unique miRNAs. Using Cytoscape [33], we visualized the circRNA-miRNA interaction network for these 612 interactions, wherein the edges between circRNAs and its target miRNAs are weighted by the number of predicted interaction sites for the circRNA-miRNA pair (Figure 2a). CDR1as was predicted to have binding sites for 74 distinct miRNAs and 63 binding sites for miR-7 (Figure 2b). According to miRTarBase [34], miR-7 interacts with 578 target genes, some of which include *SNCA* (synuclein alpha), *EIF4E* (eukaryotic translation initiation factor 4E), *KMT5A* (lysine methyltransferase 5A), *MAPKAP1* (mitogen-activated protein kinase associated protein 1), and *MKNK1* (MAP kinase interacting serine/threonine kinase 1).

**Figure. 2:**
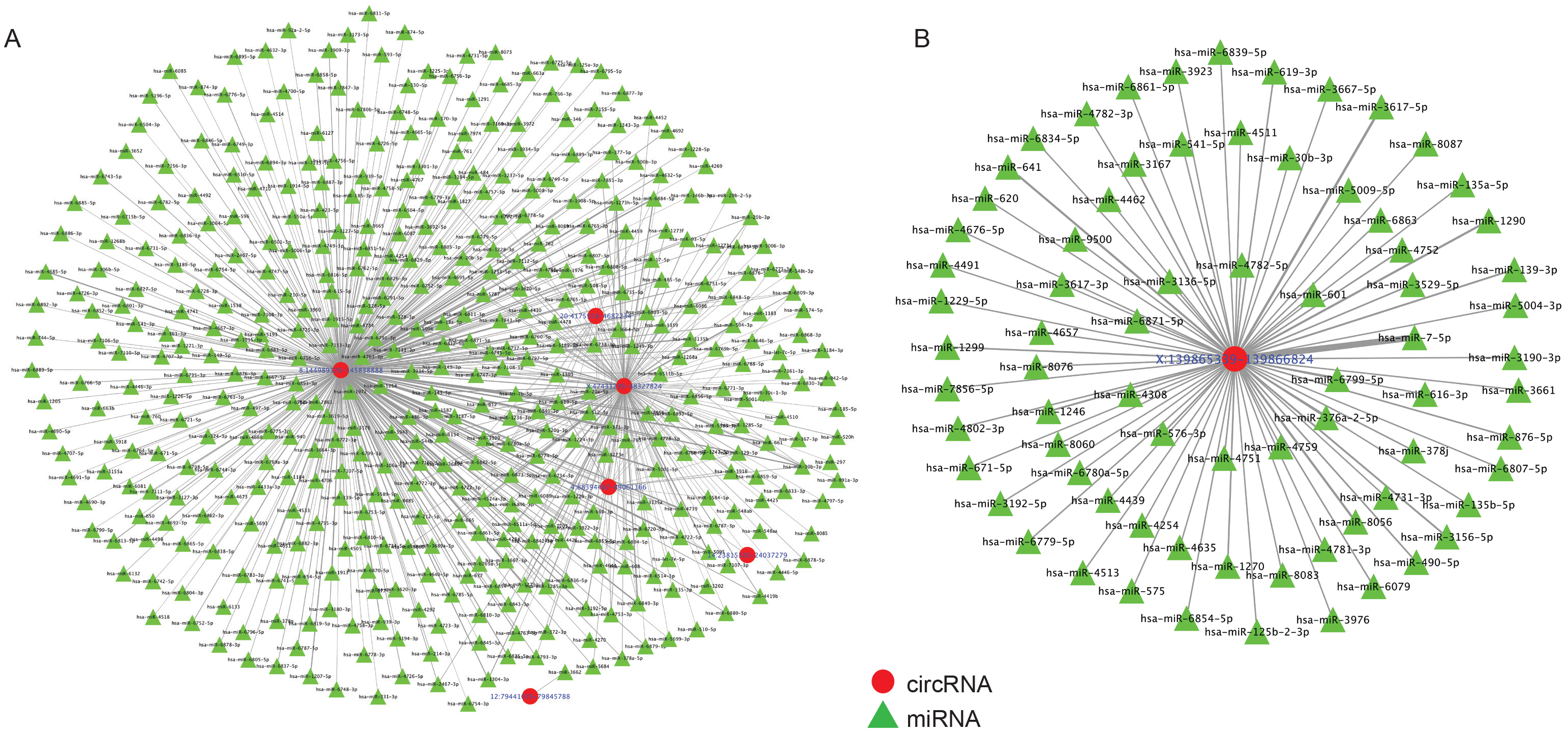
circRNA-miRNA network. (a) circRNA-miRNA interactions with 100 or more predicted binding sites. Red circular nodes: circRNAs, green triangular nodes: miRNAs. (b) miRNA network of CDR1as. The edge thickness in a and b is weighted by the number of binding sites predicted for the circRNA-miRNA interaction. miRNA, micro RNA.

We further employed the list of miRNA-mRNA target interactions common in both miRTarBase and TargetScan [35] databases, to determine the target genes of the above detected miRNAs. Overall, there were 2,530 target genes for our input list of 2,398 miRNAs, of which 255 were also differentially expressed between the AD and ND groups based on DESeq2 analysis [36] of the linear RNAs (uncorrected *p* < 0.05, Additional file 4: Table S4). Using this information about miRNA target mRNAs, we delineated a putative low-stringency circRNA-miRNA-mRNA network consisting of ten circRNAs, 53 miRNAs and 255 genes (Additional file 9: Figure S2). Further, we used the same list of circRNAs detected in ten or more samples by at least one of the circRNA prediction algorithms, and increased the filtering stringency criteria to include a miRanda match score >= 180. We also restricted the candidate miRNAs to those with mRNA targets showing differential gene expression (uncorrected *p* < 0.05) with a log2(fold change) ≥ 2 or ≤ −2 between the AD and ND groups. Using this strategy, we established a high-stringency circRNA-miRNA-mRNA interaction network with four circRNAs, 11 miRNAs and 49 genes (Figure 3, Table 1). Our overall workflow is outlined in Additional file 10: Figure S3.

**Figure. 3:**
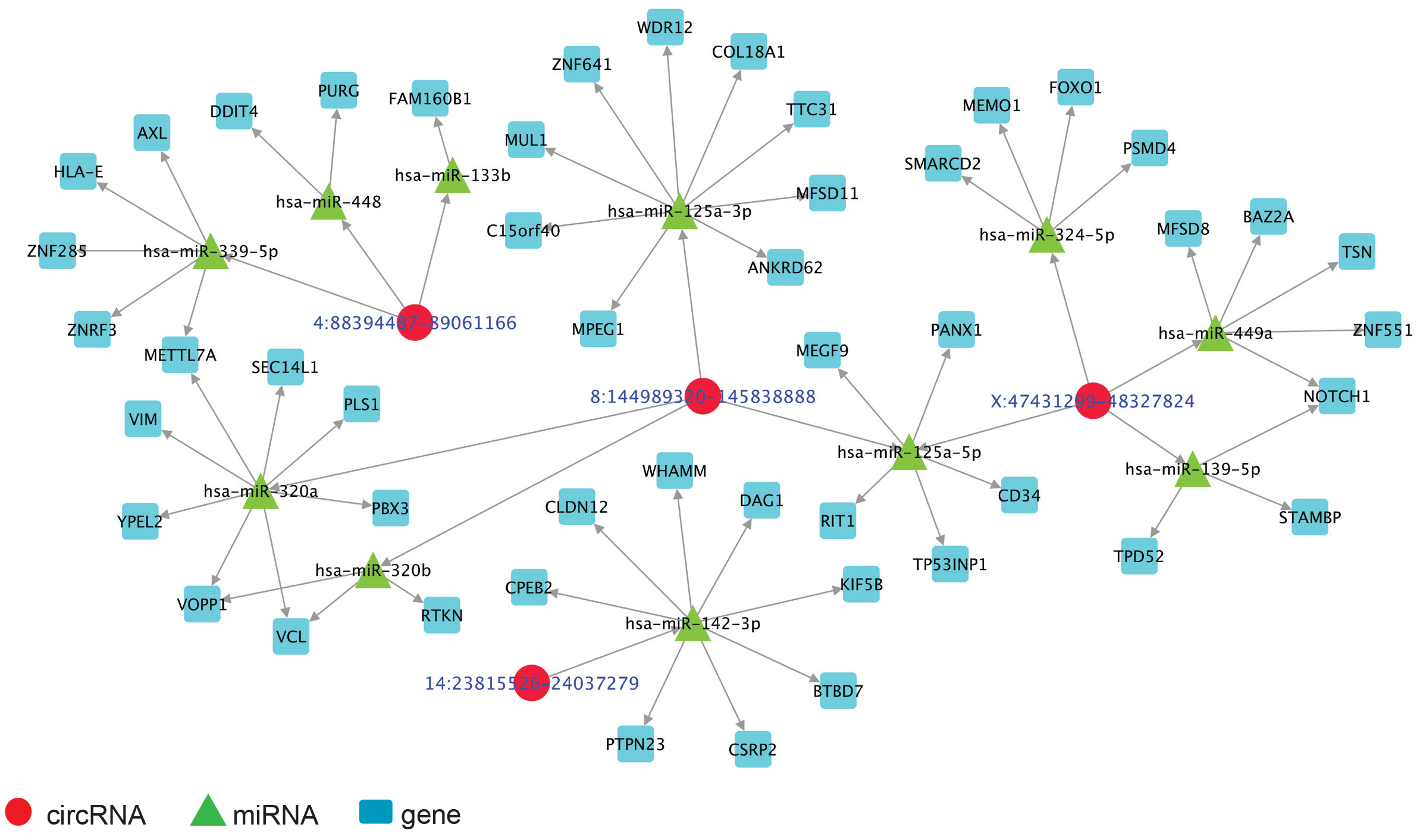
High stringency circRNA-miRNA-mRNA regulatory network. Network of circRNA-miRNA-mRNA regulation for those circRNA-miRNA interactions predicted by both RNAHybrid and miRanda, with miRanda match scores >=180 and mRNA targets with differential expression (uncorrected *p*<0.05) and log2(fold change) ≥ 2 or ≤ −2. Red circular nodes: circRNAs, green triangular nodes: miRNAs, blue square nodes: genes. mRNA, messenger RNA.

**Table 1:** circRNA-miRNA-mRNA network elements for those circRNA-miRNA interactions predicted by both miRanda and RNAHybrid, with a miRanda match score >=180and mRNA targets differentially expressed (uncorrected p < 0.05) with log2(fold change) > 2 or < −2 (high stringency network).

| Circular RNA | microRNA target | Number of binding sites predicted | Target genes (differentially expressed) |
| --- | --- | --- | --- |
| X:47431299-48327824 | hsa-miR-139-5p | 6 | NOTCH1, STAMBP, TPD52 |
| 8:144989320-145838888 | hsa-miR-320a | 2 | METTL7A, PBX3, PLS1, SEC14L1, VCL, VIM, VOPP1, YPEL2 |
| 8:144989320-145838888 | hsa-miR-320b | 2 | RTKN, VCL, VOPP1 |
| X:47431299-48327824 | hsa-miR-449a | 1 | BAZ2A, MFSD8, NOTCH1, TSN, ZNF551 |
| 8:144989320-145838888 | hsa-miR-125a-3p | 1 | ANKRD62, C15orf40, COL18A1, MFSD11, MPEG1, MUL1, TTC31, WDR12, ZNF641 |
| X:47431299-48327824 | hsa-miR-125a-5p | 1 | CD34, MEGF9, PANX1, RIT1, TP53INP1 |
| 8:144989320-145838888 | hsa-miR-125a-5p | 1 | CD34, MEGF9, PANX1, RIT1, TP53INP1 |
| X:47431299-48327824 | hsa-miR-324-5p | 1 | FOXO1, MEMO1, PSMD4, SMARCD2 |
| 14:23815526-24037279 | hsa-miR-142-3p | 1 | BTBD7, CLDN12, CPEB2, CSRP2, DAG1, KIF5B, PTPN23, WHAMM |
| 4:88394487-89061166 | hsa-miR-133b | 1 | FAM160B1 |
| 4:88394487-89061166 | hsa-miR-448 | 1 | DDIT4, PURG |
| 4:88394487-89061166 | hsa-miR-339-5p | 1 | AXL, HLA-E, METTL7A, ZNF285, ZNRF3 |

### 2.3 Pathway analysis

MetaCore pathway analysis on the 255 filtered differentially expressed target genes from the previous analysis revealed 112 perturbed pathways (corrected P < 0.01; Table 2, Additional file 5: Table S5). 23 of these were immune response-related, such as IL-4 and IL-6 signaling pathways. This identification of impacted immune response pathways is consistent with the known function of astrocytes as immune sensors in the brain and aligns with our previous RNAseq study, which showed that immune system response pathways are impacted in AD PC astrocytes compared to ND PC astrocytes [23]. Additionally,signal transduction pathways that may be perturbed include post-translational modifications (PTMs) in BAFF-induced signaling, mTORC2 downstream signaling and protein kinaseA (PKA) signaling.

**Table 2:** Top 25 pathways with FDR corrected P < 0.01 from Metacore GeneGO pathway analysis. Remaining pathways with corrected P < 0.01 summarized in Additionalfile 5: Table S5. *Ratio: Number of genes in input list found in the pathway/Total number of genes in the pathway.

| Pathway | Corrected P | *Ratio | Encoded proteins/protein complexes in input list |
| --- | --- | --- | --- |
| Aberrant B-Raf signaling in melanoma progression | 7.456E-07 | 10/55 | BAD, NOTCH1 (NICD), AKT1, Nicastrin, CBP, mTOR, ERK1/2, ERK2 (MAPK1), Mcl-1, AKT(PKB) |
| Signal transduction_PTMs in BAFF-induced signaling | 3.454E-06 | 9/51 | BAD, ERK1 (MAPK3), AKT1, MEKK1(MAP3K1), FKHR, mTOR, ERK1/2, ERK2 (MAPK1), AKT(PKB) |
| Immune response_IL-15 signaling | 9.177E-06 | 9/61 | ERK1 (MAPK3), AKT1, MEKK1(MAP3K1), mTOR, ERK1/2, UFO, ERK2 (MAPK1), Mcl-1, AKT(PKB) |
| Development_Ligand-independent activation of ESR1 and ESR2 | 9.177E-06 | 8/44 | EGFR, ERK1 (MAPK3), PKA-cat (cAMP-dependent), G-protein alpha-s, CBP, ERK1/2, ERK2 (MAPK1), AKT(PKB) |
| Translation_Regulation of EIF4F activity | 3.883E-05 | 8/54 | EGFR, MEKK1(MAP3K1), eIF4B, PP2A catalytic, eIF4A, mTOR, ERK1/2, AKT(PKB) |
| Aberrant production of IL-2 and IL-17 in SLE T cells | 5.718E-05 | 8/58 | PP2A cat (alpha), NOTCH1 (NICD), PKA-cat (cAMP-dependent), CBP, mTOR, ERK1/2, AKT(PKB), NOTCH1 precursor |
| IGF family signaling in colorectal cancer | 6.001E-05 | 8/60 | ERK1 (MAPK3), IBP, MNK2(GPRK7), mTOR, ERK1/2, ERK2 (MAPK1), AKT(PKB), Clusterin |
| Apoptosis and survival_BAD phosphorylation | 6.001E-05 | 7/42 | BAD, EGFR, PKA-cat (cAMP-dependent), PP2A catalytic, G-protein alpha-s, ERK1/2, AKT(PKB) |
| Influence of smoking on activation of EGFR signaling in lung cancer cells | 7.422E-05 | 7/44 | EGFR, ERK1 (MAPK3), PKA-cat (cAMP-dependent), MEKK1(MAP3K1), G-protein alpha-s, ERK1/2, ERK2 (MAPK1) |
| Immune response_IL-6 signaling pathway via JAK/STAT | 1.810E-04 | 8/72 | MEKK1(MAP3K1), FKHR, CCL2, CBP, mTOR, Mcl-1, AKT(PKB), gp130 |
| Signal transduction_Erk Interactions: Inhibition of Erk | 1.810E-04 | 6/34 | MKP-3, PKA-cat (cAMP-dependent), PP2A catalytic, ERK1/2, ERK2 (MAPK1), AKT(PKB) |
| Oxidative stress_ROS-mediated activation of MAPK via inhibition of phosphatases | 1.810E-04 | 6/34 | EGFR, MKP-3, SOD2, PP2A catalytic, ERK1/2, SOD1 |
| Apoptosis and survival_NGF/ TrkA PI3K-mediated signaling | 2.255E-04 | 8/77 | BAD, N-WASP, AKT1, FKHR, ILK, mTOR, ERK1/2, AKT(PKB) |
| PGE2 pathways in cancer | 2.255E-04 | 7/55 | EGFR, ERK1 (MAPK3), TGF-alpha, PKA-cat (cAMP-dependent), G-protein alpha-s, ERK2 (MAPK1), AKT(PKB) |
| Immune response_Oncostatin M signaling via MAPK | 2.275E-04 | 6/37 | ERK1 (MAPK3), MEKK1(MAP3K1), CCL2, ERK1/2, ERK2 (MAPK1), gp130 |
| Development_Beta-adrenergic receptors transactivation of EGFR | 2.275E-04 | 6/37 | EGFR, ERK1 (MAPK3), PP2A catalytic, mTOR, ERK1/2, ERK2 (MAPK1) |
| Development_Regulation of lung epithelial progenitor cell differentiation | 3.976E-04 | 6/41 | FOXP2, SMAD4, Frizzled, NOTCH1 receptor, FOXP1, O-fucose |
| Regulation of Tissue factor signaling in cancer | 4.726E-04 | 6/43 | EGFR, ERK1 (MAPK3), mTOR, ERK1/2, ERK2 (MAPK1), AKT(PKB) |
| Development_Adenosine A2A receptor signaling | 4.726E-04 | 6/43 | BAD, PKA-cat (cAMP-dependent), MEKK1(MAP3K1), G-protein alpha-s, ERK1/2, AKT(PKB) |
| Development_VEGF signaling via VEGFR2 - generic cascades | 5.478E-04 | 8/93 | ERK1 (MAPK3), MEKK1(MAP3K1), CCL2, Vinculin, ERK1/2, ERK2 (MAPK1), AKT(PKB), Fyn |
| Signal transduction_mTORC2 downstream signaling | 5.478E-04 | 7/68 | BAD, AKT1, PKA-cat (cAMP-dependent), FKHR, mTOR, Mcl-1, AKT(PKB) |
| Transcription_Hypoxia- and receptor-mediated HIF-1 activation | 5.478E-04 | 6/46 | EGFR, FKHR, MNK2(GPRK7), CBP, ERK1/2, AKT(PKB) |
| Transcription_Androgen Receptor nuclear signaling | 5.478E-04 | 6/46 | EGFR, AKT1, PKA-cat (cAMP-dependent), Frizzled, ERK2 (MAPK1), AKT(PKB) |
| Signal transduction_PTEN pathway | 5.478E-04 | 6/46 | BAD, EGFR, ILK, mTOR, ERK1/2, AKT(PKB) |
| Immune response_IL-4 signaling pathway | 5.478E-04 | 8/94 | BAD, PDE3A, MEKK1(MAP3K1), FKHR, mTOR, ERK1/2, ERK2 (MAPK1), AKT(PKB) |

### 2.4 Lack of circRNA differential expression in AD PC astrocytes

We analyzed our catalog of circRNA candidates to determine whether there were circRNAs uniquely expressed in either the AD or ND cohort. Though there were over 2,000 circRNAs unique to each group, we did not observe them to be recurrent in the samples within their respective group. The log2 (fold change) for all candidates calculated using DESeq2 are summarized in Additional file 1: Table S1. 93 circRNAs were unique to AD and called in at least two samples by at least one of the tools, and 82 circRNAs were unique to ND and called in at least two samples by at least one of the tools. These circRNA candidates were supported by at least two junction reads. To identify any differentially expressed candidates, we performed a Student’s t-test on those circRNAs commonly called across the two groups. Only two circRNAs trending towards significance (uncorrected *p* < 0.05) were identified and include 1:201452657-201736927 (uncorrected *p* = 0.015) and 16:1583657-2204141 (uncorrected *p* = 0.046).

## 3. Discussion

CircRNAs, which are abundant in the mammalian brain, represent a recent addition to the class of non-coding RNAs. In this study, we detected astrocytic circRNAs using whole transcriptome RNAseq data obtained from the PC of AD and ND subjects, and outlined circRNA-miRNA-mRNA regulatory networks. Based on the results from four different circRNA detection algorithms, we identified over 4000 unique circRNAs across all samples, the majority of which were derived from coding exons. Although we did not identify circRNAs that were differentially expressed and recurrent in AD or ND, we were able to delineate circRNA-miRNA-mRNA networks for the ten most recurrent circRNAs expressed across both groups, and also incorporate our previous differential expression analysis data from the linear mRNA. We observe that the majority of identified circRNAs are unique in the AD or ND groups and are not recurrent across the respective groups. This could be due to their low abundance in those samples, which may be below detection levels, or could be due to biological differences between the two groups, which requires further investigation. Pathway analysis on the differentially expressed miRNA target genes identified immune system related and signal transduction pathways. Notably, astrocytes are active players in cerebral innate immunity [37], and previous studies have reported that astrocytes respond to IL-4 signaling and potentially mediate between the immune effector cells and the nervous responders[38]. These predicted regulatory network and pathway analyses may help provide new insights into transcriptional regulation in the brain.

The circRNA CDR1as (also known as CiRS-7, a circRNA sponge for miR-7) was detected in all 20 of our samples and is a widely reported circRNA with 63 conserved seed matches for miR-7, indicating possible miR-7 binding sites [3, 11]. Interestingly, overexpression of CDR1as in zebrafish decreased the midbrain size, suggesting a functional role for CDR1as in the brain, while a knock down of CDR1as downregulated miR-7 targets in HEK293 cells [3]. This regulation is relevant since miR-7 plays a role in Parkinson’s disease, stress handling and brain development [3, 39], and also has tumor-suppressive properties [39]. CDR1as also showed widespread expression in neuroblastoma and astrocytoma [40]. However, the expression of CDR1as was reduced in AD hippocampal samples about 0.18-fold compared to controls [15], which we did not observe in our PC astrocyte dataset. Apart from CDR1as,the tools also predicted circRNAs derived from genes such as *SLC8A1* (solute carrier family 8 - sodium/calcium exchanger - member 1), which is under-expressed in hippocampal neurons from aged human brains [41], *SYT1* (synaptotagmin 1), whose increase was correlated to age-related spatial cognitive impairment in mice [42], *PSAP* (prosaposin), which is increased in activated glia during normal aging in mouse brains [43], and *FGF17* (fibroblast growth factor 17).

Although our dataset provides insights into the existence and abundance of astrocytic circRNAs in elderly individuals, there are a few limitations. Primarily, the whole-transcriptome data we analyzed was not generated from samples that were depleted of linear RNAs using RNase R (ribonuclease R), an exoribonuclease that selectively digests linear RNA but leaves behind lariat or circRNA structures. Due to the presence of a larger pool of transcripts, which are mostly linear RNAs, RNAseq may not have comprehensively captured all the circRNAs in the samples. Notably, this enrichment step has been used by various groups to enrich for circRNAs for sequencing analyses [2, 3, 44].

Another limitation of bioinformatics-based circRNA detection is the highly divergent results produced by different algorithms. We observed this in our analyses and it has also been reported by two recent circRNA benchmark studies [29, 45]. The algorithms utilize different aligners, heuristics and filtering criteria, thus introducing ‘blind spots’ (false negatives) when addressing biases introduced by each method [46]. For example, find_circ and CIRI rely on filtering for GT-AG splice signals and thus may not capture candidates with non-canonical splice signals. Further, most tools use a read count filter, which may not be ideal for circRNAs with low expression relative to their linear host [47]. Given the low reliability on read counts, statistical approaches improve detection and classification of splice junctions, including novel ones [48]. Among the circRNA detection algorithms, KNIFE implements a logistic generalized linear model to distinguish true circRNAs, and is therefore able to identify circRNAs derived from non-canonical splice sites. Notably, KNIFE achieves a more balanced performance, for precision and sensitivity, compared to other circRNA detection algorithms, as described in one of the benchmarking studies [45]. We observed in out dataset that KNIFE detected more circRNAs compared to find_circ, CIRI and CIRCexplorer. Nonetheless, sequencing errors and technical artifacts introduced during RNAseq can still lead to false positive circRNAs, and hence statistical tests to estimate false discovery rates in circRNA detection need to be developed.

While circRNAs have continued to gain attention as an abundant non-coding RNA species with potential regulatory functions, our understanding of their expression in various cell and tissue types remains limited. To address this challenge, we describe an analysis of astrocytic circRNAs in RNAseq data from elderly individuals, and we delineate potential circRNA-miRNA-mRNA regulatory networks. Given the role of astrocytes in signaling and synaptic modulation, and as immune sensors in the brain, the circRNAs we identified may be associated with such key functions. Further characterization using circRNA-enriched datasets will help us understand the atlas of circRNA expression in the context of specific cell types and conditions, including aging and AD. In addition, downstream functional studies are needed to clarify how and whether circRNAs act as hubs for influencing protein expression and cellular processes. As we continue to piece together the factors involved in transcriptional regulation, we will both better understand basic cellular mechanisms and set the stage for developing improved therapeutic strategies for AD and other diseases.

## 4. Conclusions

In summary, we utilized astrocyte specific RNAseq data to identify astrocytic circRNAs in aged subjects (N=20). Utilizing four circRNA prediction algorithms, we identified a total of 4,438 unique circRNAs across samples, majority of which were derived from exonic regions. The widely reported CDR1as circRNA was detected in all 20 samples with a median of 52 supporting reads. Given the putative miRNA regulatory function of circRNAs, we further performed an in silico prediction of putative miRNA binding sites on the ten most recurrent circRNAs, and further delineated a low- and high-stringency circRNA-miRNA-mRNA regulatory network. Pathway analysis on the genes from our low-stringency network revealed significantly impacted immune response pathways, which aligns with the known function of astrocytes as immune sensors in the brain. While we did not detect circRNAs recurrently expressed in the context of healthy controls or Alzheimer’s, we are the first to report circRNAs and their potential regulatory impact in a cell-specific and region-specific manner in aged subjects. Continued analyses such as these sets the foundation for circRNA characterization and understanding their expression and regulatory networks in specific cell types and regions in the brain.

## 5. Methods

### 5.1 Sample acquisition, library preparation and paired-end sequencing

Detailed methods for sample acquisition, immunohistochemistry using an aldehyde dehydrogenase 1 family, member L1 (ALDH1L1) antibody, microdissection, RNAseq library preparation and sequencing of astrocytes are described in our previous publication [23]. Briefly, postmortem human brain samples were collected at the Banner Sun Health Research Institute’s (BSHRI) Brain and Body Donation Program (BBDP) from 10 clinically classified LOAD subjects (4 males and 6 females; 5 APOEε3/4 subjects and 5 APOEε3/3 subjects) and 10 ND controls (6 males and 4 females; 5 APOEε3/4 subjects and 5 APOEε3/3 subjects). All subjects were enrolled in the BSHRI BBDP in Sun City, Arizona, and written informed consent for all aspects of the program, including tissue sharing, was obtained either from the subjects themselves prior to death or from their legally-appointed representative. The protocol and consent for the BBDP was approved by the Western Institutional Review Board (Puyallap, Washington). Clinical and pathological donor demographics are summarized in Additional file 6: Table S6. Approximately 300 astrocytes were laser capture microdissected from PC brain sections and total RNA was isolated from the cell lysates, followed by cDNA creation and library generation. Equimolar pools of libraries were sequenced by synthesis on the Illumina HiSeq2000 for paired 83 base pair reads.

### 5.2 Data analysis

The data analysis workflow is summarized in Additional file 10: Figure S3. Raw sequencing data, in the form of basecall files (BCLs), were converted to FASTQ format using Illumina’s bcl2fastq conversion software and quality checked using FastQC [49]. To eliminate variance in circRNA detection that could arise due to differences in the number of sequencing reads, all FASTQ files were down-sampled to 85,547,262 reads using seqtk [50]. The down-sampled FASTQ files were then run through four different circRNA prediction algorithms—CIRCexplorer (v1.1.10), CIRI (v2), find_circ (v1), and KNIFE (v1.4), using the parameter settings described in Additional file 7: Table S7. CircRNAs from each sample with at least two supporting reads were used for further downstream processing and analyses. CIRI produces 1-based circRNA coordinates, and was therefore converted to 0-based coordinates to be consistent with the other three algorithms. We then annotated our catalog of circRNA candidates using the UCSC RefSeq annotations [51] and BEDtools [52].

The ratio of circular to linear RNA isoforms was calculated using the approach described in [8]. For each circRNA candidate, we used the number of back-spliced reads for circRNA quantification (*N_c_*) and the number of linear reads supporting the same 5’ or 3’ splice junction (*N*_*l5*_ or *N*_*l3*_) as the number of linear RNA reads. The linear junction supporting reads were obtained by aligning our RNAseq data to the reference genome (GRCh37) using STAR [53].

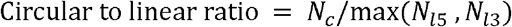

#### 5.2.1 miRNA target prediction

For circRNAs detected in at least 50% of the samples, we next conducted miRNA binding site prediction using the miRanda [30] and RNAHybrid [31] algorithms. The miRanda algorithm finds potential target sites for miRNAs in a genomic sequence using a two-step strategy. First, a dynamic programming local alignment is implemented between the miRNA sequence and the sequence of interest (circRNA sequence in this study), scoring the alignment based on sequence complementarity (match score). In the second step, the thermodynamic stability of the resulting RNA duplex is estimated based on the high-scoring alignments from the first phase. The RNAHybrid algorithm finds the energetically most favorable hybridizations of a small RNA to a large RNA. Only those circRNA-miRNA interactions predicted by both the algorithms are used for our downstream network construction and analyses. From the list of commonly predicted circRNA-miRNA interactions, we filtered for those having a miRanda match score >=150.

#### 5.2.2 circRNA-miRNA-mRNA network construction

miRNA-mRNA interactions that are common in both miRTarBase [34] and TargetScan [35] were then used to determine the gene targets of each filtered miRNA and compared with genes identified using differential expression analysis of the linear RNAs (uncorrected *p* < 0.05; DESeq2 performed as described in our previous publication). Using these data, we outlined a low-stringency circRNA-miRNA-mRNA regulatory network with custom python scripts and visualized the network using cytoscape. We further filtered for the circRNA-miRNA interactions with miRanda match scores >=180 and miRNAs with mRNA targets showing differential expression (uncorrected *p* < 0.05, log2[fold change] ≥ 2 or ≤ −2) to outline a high-stringency circRNA-miRNA-mRNA network.

#### 5.2.3 Pathway analysis

On the list of filtered miRNA target genes with DESeq2 uncorrected *p* < 0.05, we performed pathway analysis using MetaCore GeneGO (v6.32.69020) from Thompson Reuters to predict pathways that are commonly impacted in the AD and ND groups. The results were filtered for enriched pathways with a false discovery rate (FDR)-corrected P < 0.01.

## Abbreviations

CircRNA: circular RNA
miRNA: microRNA
RNAseq: RNA-sequencing
PC: posterior cingulate
AD: Alzheimer’s disease
LOAD: Late-onset Alzheimer’s disease
CDS: coding DNA sequences
mRNA: messenger RNA
FDR: False discovery rate
ALDH1L1: aldehyde dehydrogenase 1 family, member L1
BSHRI: Banner Sun Health Research Institute
BBDP: Brain and Body Donation Program

## Declarations

### Ethics approval and consent to participate

All subjects were enrolled in the BSHRI BBDP in Sun City, Arizona, and written informed consent for all aspects of the program, including tissue sharing, was obtained either from the subjects themselves prior to death or from their legally-appointed representative. The protocol and consent for the BBDP was approved by the Western Institutional Review Board (Puyallap, Washington).

### Consent for publication

Not applicable

### Availability of data and materials

All the RNAseq data generated in this study are accessible through the National Center for Biotechnology Information (NCBI) database of Genotypes and Phenotypes (dbGaP; accession# phs000745.v1.p1), and data supporting our findings are included within the manuscript and additional figures/tables.

### Competing interests

The authors declare that they have no competing interests

### Funding

Research reported in this publication was supported by the National Institute on Aging (NIA) of the National Institutes of Health under award number P30AG019610, and the Arizona Department of Health Services award number ADHS14-052688. The content is solely the responsibility of the authors and does not necessarily represent the official views of the National Institutes of Health. The funders had no role in the study design, data collection and analysis, decision to publish, or preparation of the manuscript.

### Authors’ contributions

SS and WL conceived the study and wrote the manuscript. SS performed all the data analysis and interpretation. LC, JA and PG made substantial contribution to data acquisition, performed library preparation and sequencing of all samples and also contributed to data interpretation. All authors read and approved the final manuscript.

## Acknowledgements

We are grateful to the Banner Sun Health Research Institute Brain and Body Donation Program of Sun City, Arizona for the provision of human brain tissues. The BBDP has been supported by the National Institute of Neurological Disorders and Stroke (U24 NS072026 National Brain and Tissue Resource for Parkinson’s Disease and Related Disorders), the National Institute on Aging (P30AG19610 Arizona Alzheimer’s Disease Core Center), the Arizona Department of Health Services (contract 211002, Arizona Alzheimer’s Research Center), the Arizona Biomedical Research Commission (contracts 4001, 0011, 05-901 and 1001 to the Arizona Parkinson’s Disease Consortium) and the Michael J. Fox Foundation for Parkinson’s Research [54]. We would also like to thank TGen’s Dr. Kendall Jensen and Dr. Elizabeth Hutchins for input and guidance, and Cynthia Lechuga for administrative support. Nancy Linford, PhD, provided editorial suggestions.

## Additional files

Additional file 1: Table S1. Summary of all detected circRNAs. (.xlsx)

Additional file 2: Table S2. Circular to linear ratios for all detected circRNAs

Additional file 3: Table S3. circRNA-miRNA interactions with ≥ 100 predicted binding sites. (.xlsx)

Additional file 4: Table S4. DESeq2 analysis results for genes with uncorrected p < 0.05, between AD and controls. (.xlsx)

Additional file 5: Table S5. Pathways with corrected P < 0.01, apart from the ones summarized in Table 2. (.xlsx)

Additional file 6: Table S6. Donor demographics. (.xlsx)

Additional file 7: Table S7. Tool parameters used for circRNA detection in this study. (.xlsx)

Additional file 8: Figure S1. Circular to linear ratios for all detected circRNAs. (.pdf)

Additional file 9: Figure S2. Low stringency circRNA-miRNA-mRNA regulatory network. (.pdf)

Additional file 10: Figure S3. Computational workflow outline and filtering criterion. (.pdf)

**Figure S1:** Circular-to-linear ratios. Ratio of average back-spliced reads to average linearly spliced reads for all detected circRNAs.

**Figure. S2:** Low stringency circRNA-miRNA-mRNA regulatory network. Network of circRNA-miRNA-mRNA regulation for those circRNA-miRNA interactions predicted by both RNAHybrid and miRanda, with miRanda match scores >= 150 and mRNA targets with differential expression (uncorrected *p* < 0.05). Red circular nodes: circRNAs, green triangular nodes: miRNAs, blue square nodes: genes

**Figure. S3:** Computational workflow outline and filtering criterion. PC, posterior cingulate; RNAseq, RNA sequencing; circRNA, circular RNA; miRNA, micro RNA; mRNA, messenger RNA.

